# Amniotic fluid extracellular vesicle properties evolve with gestational age and reflect fetal development

**DOI:** 10.1101/2025.04.23.649052

**Authors:** Ishara Atukorala, Sally Beard, Ching-Seng Ang, Hamish Brown, Swetha Raghavan, Natasha de Alwis, Bianca Fato, Natalie Binder, Natalie Hannan, Lisa Hui

## Abstract

Amniotic fluid (AF) is a valuable source of extracellular vesicles (EVs) derived from the fetoplacental unit. Preclinical and clinical studies have highlighted promising applications of AF-EVs and their role in cellular communication, yet our understanding of AF-EV physiology is limited. This study aimed to examine the physiological importance of AF-EVs in fetal development from the second trimester to term gestation. We obtained AF samples from routine second-trimester amniocentesis and prelabour Caesarean section at term. We isolated EVs using a combination of differential centrifugation, filtration, and ultracentrifugation and characterised them using nanoparticle tracking analysis, cryoelectron microscopy, and Western blotting. The differential EV proteome was analysed using label-free proteomics. We assessed the second trimester and term AF-EV properties through an enrichment analysis. The EV size and protein enrichment difference revealed a gestational-age-dependent variation in the predominant EV subtype. Second-trimester-derived EVs were enriched in ectosomes, while term EVs contained a significant proportion of exosomes. We identified several morphologies of AF-EVs, including unilamellar, multilamellar, multicompartmental and granular-centred EVs, across gestations. Proteomics analysis of AF-EVs identified 4137 proteins with high confidence, of which 1099 exhibited significant differential expression between the two groups. Second-trimester-enriched AF-EV proteins represented molecule assembly processes, metabolism and organogenesis. At term, AF-EV proteins corresponded to impending newborn functions such as immunity and digestion. In conclusion, we provide compelling evidence that EV biogenesis and secretion in the fetoplacental unit undergo significant alterations across gestation, revealing a complex and dynamic physiology and intercellular communication that adapts to the needs of the developing fetus.

Graphical abstract
This figure summarises the study’s main findings, including the shift in the predominant EV subtype, the different organs and biofluids represented by the enriched proteins at each gestation, and the main biological pathways.

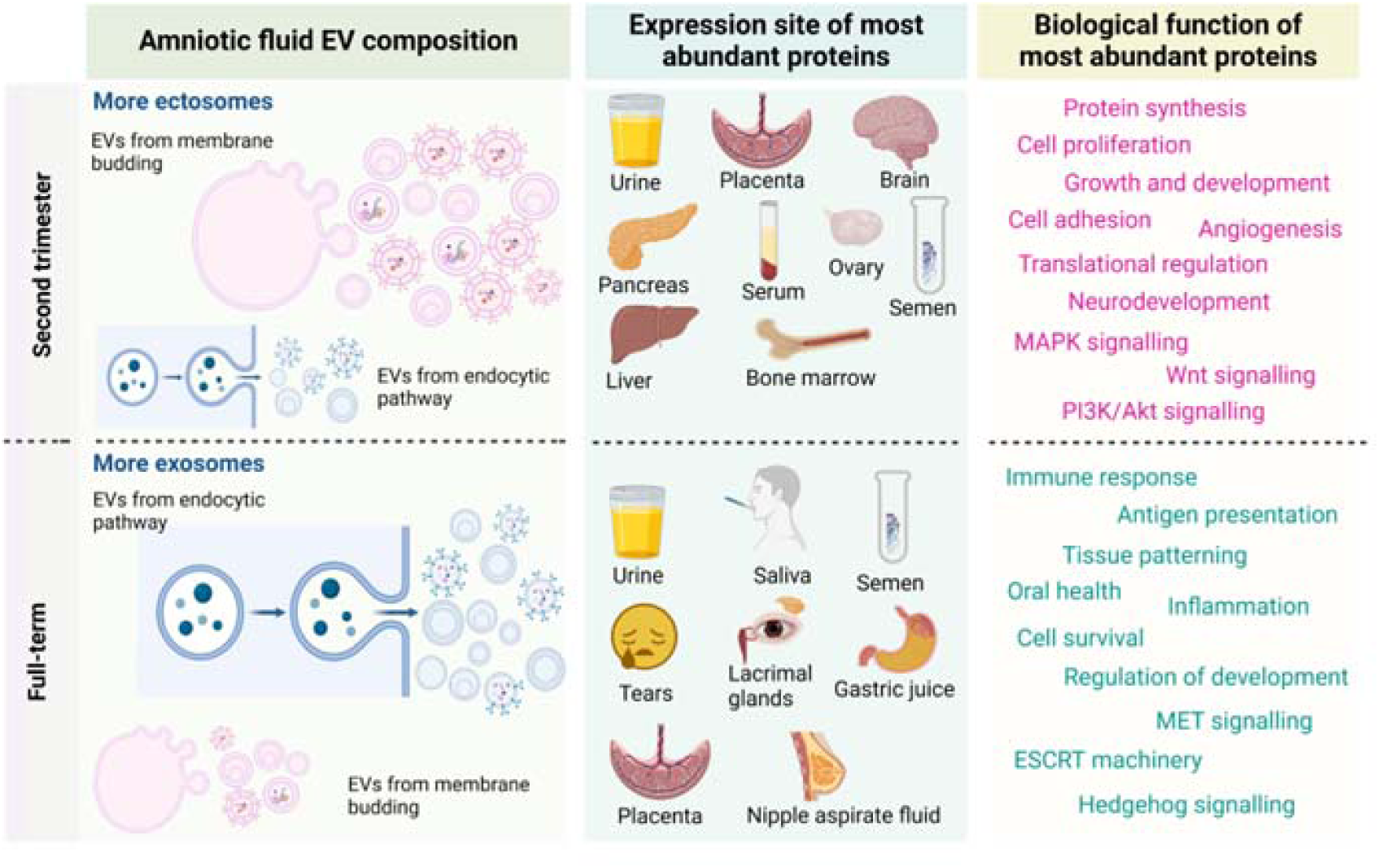

## Background

Extracellular vesicles (EVs) are released from all living cells as a natural aspect of their cellular homeostasis. They are membrane-enclosed vesicles that contain selectively packed cargo of biologically active molecules such as proteins, nucleic acids and lipids (1, 2). EVs mediate intercellular communication via this internal cargo and membrane-bound molecules. Their role in facilitating inter-organ communication, coordinating physiological processes and maintaining systemic homeostasis is widely studied (3–5). EV biogenesis pathways influence cargo sorting (6) and also change in response to extracellular stimuli (7, 8). However, the mechanisms driving the change in EV cargo composition, particularly *in vivo*, are poorly understood (9).

Amniotic fluid (AF) is a valuable sample with a wealth of fetal biological information (10–12). It is used clinically for prenatal diagnosis, primarily for amniocyte analysis for chromosome and genetic abnormalities (13, 14). However, AF-EVs have been relatively unexplored until recently. Numerous exploratory studies have been published in the growing field of AF-EV research (15), but relatively little attention has been paid to the basic phenotypic and physiological properties of AF-EVs across gestation.

EV biology in pregnancy is an emerging area of research, with substantial potential for translational impact in prognosis and treatment in perinatal medicine. The lipid bi-layer membrane of EVs protects their cargo from external degrading enzymes (16), suggesting that EVs can contain stable biomarkers for understanding health and disease (17, 18). A growing body of pre-clinical and clinical researchers are now investigating the anti-inflammatory, regenerative and angiogenic molecules within EVs for therapeutic applications (19, 20). However, our understanding of the general physiology of EVs from the fetoplacental unit is limited, including the cells they represent, the molecular information contained in them, and the likely purpose of this information for the baby and the mother.

This study aimed to delineate and compare AF-EVs derived from uncomplicated second-trimester and term pregnancies to uncover their physiology. These findings may enhance our understanding of AF-EV biogenesis, cargo, and physiological significance in fetal growth and development.

## Methods

### Amniotic fluid sample collection

AF samples were collected prospectively at Mercy Hospital for Women in Melbourne, Australia, following written informed participant consent. Second-trimester samples were collected from clinically-indicated amniocentesis due to abnormal prenatal screening or sonography results (Table 1). Term samples were collected from pre-labour Caesarean sections after the uterine incision prior to membrane rupture, as previously described (21)The AF samples were centrifuged at 300 x g for 10 minutes to remove cells and debris before being stored at −80oC until EV isolation. This study was approved by the Human Research Ethics Committee of Mercy Health (2023–014).

**Table 1.** Clinical characteristics of the samples.

| Second-trimester samples |  |  | Full-term samples |  |
| --- | --- | --- | --- | --- |
| Gestation at collection (weeks) | Fetal sex | Indication for amniocentesis | Gestation at collection (weeks) | Fetal sex |
| 15.4 | F | Bilateral cleft lip and palate | 38.2 | M |
| 15.4 | M | Thin nuchal translucency (4.5mm) | 39.0 | F |
| 16.1 | M | Increased chance of trisomy 21 detected in non-invasive prenatal testing | 39.0 | F |
| *16.2 | F | Increased chance of trisomy 21 detected in first trimester combined screening | 39.3 | M |
| 23.0 | M | Bilateral mild cerebral ventriculomegaly | 39.0 | M |
| 23.1 | F | Increased chance of trisomy 21 detected in maternal serum screening | 39.3 | F |

*F, Female; M, Male*
*\*Birth via Caesarean section at 36 weeks due to placenta praevia. Other women in the second-trimester amniocentesis group delivered healthy live newborns at term gestation.*

### EV isolation

AF-EVs were isolated using differential centrifugation, filtration, and ultracentrifugation. EVs were characterised according to MISEV23 guidelines (22).

AF samples were thawed overnight at 4°C and centrifuged at 5000 g for 20 minutes at 4°C to remove vernix caseosa and cell debris. Supernatant was centrifuged to 15,000 g for 30 minutes at 4°C, then filtered through 0.22 µm filter. The resulting supernatant was subjected to ultracentrifugation at 120,000 g for 2 hours at 4°C to precipitate small EVs. EV pellet was then washed with 0.22 μm-filtered Dulbecco’s phosphate-buffered saline (DPBS; Gibco™) and centrifuged at 120,000 g for 2 hours at 4°C. The resulting EV pellet was resuspended in DPBS and stored at −80°C until further analysis.

### EV protein quantification

EV proteins (2 µg) of each sample immobilised on a polyacrylamide gel was fixed with fixer buffer (50% Methanol, 7% Acetic acid) and stained with Sypro Ruby gel stain (Thermo Fisher Scientific) for 24 hours. After washing the gel with wash buffer (10% methanol, 7% acetic acid), the gel was scanned using Chemidoc (BioRad) and the lanes were quantified against Benchmark^TM^ protein ladder (Thermo Fisher Scientific).

### Nanoparticle Tracking Analysis

EV size range and concentration was determined using NanoSight NS300 (Malvern Panalytical; NanoSight NTA 3.2 software). Samples were diluted 400-fold with 0.22 μm-filtered PBS and injected at an infusion rate of 50. Each sample was captured in 3 rounds of 30 seconds, 3 times with camera level and detection threshold set at 11 and 5, respectively, at 25°C.

### Western blotting

EVs corresponding to 15 µg of protein were lysed in 4x Laemmli buffer (8% (w/v) SDS, 10% (v/v) glycerol, 200mM Tris-HCL pH 6.8 and trace of bromophenol blue), and 2M DTT, heated at 95°C for 2 min and separated on 4-15% polyacrylamide gels (BioRad). Gels were electrophoresed at 120 V for ∼90 min. Proteins were transferred to PVDF membranes (Thermo Fisher Scientific) and blocked with 5% skim milk. Membranes were incubated with primary antibodies for Alix (Catalog # E6P9B) (1:1000) (Cell signalling), CD9 (Catalog # 10626D) (1:500) (Thermo Fisher Scientific), CD63 (Catalog # 10628D) (1:500) (Thermo Fisher Scientific), at 4°C overnight. After washing the excess antibody off, the membranes were incubated with the relevant fluorescent-conjugated secondary antibodies (IRDye 680RD Goat anti-Rabbit IgG or IRDye 800CW Goat anti-Mouse IgG) (LICORbio) in 1:10,000 dilution, for 1 h at room temperature. Membranes were imaged using ChemiDoc (BioRad).

### Cryo-electron microscopy and image analysis

Cryo-electron microscopy was used to visualise vesicles. Gold 300 mesh, lacey carbon film coated EM grids (ProSciTech, Australia) were glow-discharged (15 mA, 30 sec) using GloQube® Plus Glow Discharge System. Sample (3 μL of initial EV preparation) was applied on to the carbon side of the grid, which was then blotted for 4.0 s (−1 bolt force) with Whatman filter paper #1 and plunge-frozen into liquid ethane using a Thermo Fisher Vitrobot Mark IV. The climate chamber was maintained at 90% humidity, 4°C. EM grids containing frozen samples were stored in liquid nitrogen until imaging. EVs were visualised using a Thermo Fisher TECNAI F30 cryo-electron microscope, equipped with a Ceta electron camera, at the Bio21 Advanced Microscopy Facility (The University of Melbourne). Images were recorded at x2400 magnification with a defocus of −5 μm and 10 1 pixel size.

Image analysis tool Fiji ImageJ (23) was used to measure the diameter of EVs. First, the scale bar of each image was defined by the number of pixels under the “set scale” function. Then the EVs were measured in 2 perpendicular measurements. Since the average measurement would misinterpret the non-circular morphology of EVs, the larger measurement was used for the analysis.

### Sample preparation for Mass spectrometry

EV proteins (30 µg) were lysed in 2X lysis buffer (10% SDS, 100 mM TEAB pH 8.5). Disulphide bonds of the proteins were reduced using Tris-(2-carboxyethyl)phosphine (final concentration 5 mM) (Thermo Fisher Scientific) and alkylated using Methyl methanethiosulfonate (final concentration 20 mM). Samples were acidified with a final concentration ∼2.5% phosphoric acid. The samples were loaded onto S-Trap™ micro spin columns (ProtiFi, USA) with binding/wash buffer (100 mM triethylammonium bicarbonate in 90% methanol) and centrifuged at 4,000 g for 30 seconds to trap proteins. After washing the columns 3 times with 150 μL binding/wash buffer, proteins were digested with Sequencing Grade Modified Trypsin (Promega) at a ratio of 1:10 (w/w) trypsin: protein for 2 hours at 47°C. Peptides were collected consecutively in 3 elution steps with 50 mM TEAB in water, 0.2% formic acid in water, and 50% acetonitrile in water. Eluted peptides were lyophilised using a vacuum Concentrator (Savant SpeedVac™) and reconstituted in mass spectrometry sample buffer (2% acetonitrile, 0.05% trifluoroacetic acid) to a final peptide concentration of 0.5 µg/µL.

### Data-independent acquisition (DIA) Mass spectrometry (MS)

Liquid chromatography (LC) based tandem mass spectrometry (MS/MS) was carried out using an Orbitrap Ascend mass spectrometer (Thermo Fisher Scientific) equipped with a nanoflow reversed-phase-HPLC (Ultimate 3000 RSLC, Dionex), fitted with an Acclaim Pepmap nano-trap column (Dionex—C18, 100 Å, 75 µm× 2 cm) and an Acclaim Pepmap RSLC analytical column (Dionex-C18, 100 Å, 75 µm× 50 cm), at the Melbourne Mass Spectrometry and Proteomics Facility (The University of Melbourne). The tryptic peptides were injected into the enrichment column at an isocratic flow of 5 µL/min of 2% v/v CH_3_CN containing 0.1% v/v formic acid, for 5 min before the enrichment column was switched in line with the analytical column. The eluents for LC were 5% DMSO in 0.1% v/v formic acid (solvent A) and 5% DMSO in 100% v/v CH_3_CN and 0.1% v/v formic acid (solvent B). The flow gradient was, i) 3% B for 0-6 min, ii) 3-4% B for 6-7min, (ii) 4-25% B for 7-82 min, (iii) 25-40% B for 82-86min, (iv) 40-80% B for 86-87min, (v) 80-80% B for 87-90min and (vi) 80-3% for 90-91min. The column was equilibrated at 3% B for 10 minutes before the next sample injection.

For DIA experiments, full MS resolutions were set to 120,000 at m/z 200 and scanning from 350-1400 m/z in the profile mode. Full MS Automatic Gain Control target was 250% with an injection time (IT) of 50 ms. AGC target value for fragment spectra was set at 2000%. Fifty windows of 13.7 Da were used with an overlap of 1 Da. Resolution was set to 30,000 and maximum IT to 55 ms. The normalised collision energy was set at 30%. All data were acquired in centroid mode using positive polarity.

### Proteomics data analysis

DIA data were analysed using the direct DIA analysis workflow with default settings on the Spectronaut® software (v. 17.5.230413.55965) and against the UniProt Homo Sapiens database (updated Sep 2023). Trypsin specificity was set to two missed cleavages.

Carbamidomethyl (Cys) was defined as the fixed modification, while acetylation (protein N-term) and oxidation (Met) were defined as variable modifications. Results were filtered at a protein and peptide spectrum matched with a false discovery rate of 1%. Precursor filtering used the Q value, and quantification was done at the MS2 level. The cross-run normalisation strategy was set to automatic. Quantitative data were exported for statistical analyses onto MaxQuant Perseus (Max-Planck-Institute of Biochemistry) (24). The data was log2 transformed and annotated to the 2 groups: second trimester and full-term. The rows were filtered to include the proteins identified in all 12 samples.

Ubiquitously expressed peptides in all samples were subjected to the Student’s T-test (with a false discovery rate of 0.05 and S0 filter set to 0.1) to determine significantly differentially abundant proteins between groups. FunRich Version 3.1.4 was used to perform the enrichment analysis. It is a functional enrichment analysis tool that integrates multiple databases including Uniprot, Human Protein Reference Database (HPRD) and Human Protein Atlas (25).

### Statistical analysis

The statistical analysis for all experiments except proteomics was conducted using GraphPad Prism 10.1.0. We determined the normal distribution of the data using the Shapiro-Wilk test. Unpaired two-tailed Student’s T-test and the Mann-Whitney U test were used for parametric and nonparametric datasets, respectively.

## Results

### Clinical characteristics of the samples

Twelve AF samples were analysed; six from second-trimester amniocenteses and six from term prelabor Caesarean sections. Each group comprised three female and three male fetuses. Of the second trimester group, three underwent amniocentesis for an increased chance of trisomy 21 on prenatal screening and others due to fetal ultrasound anomalies. One 23-week male fetus with mild bilateral ventriculomegaly had no additional findings on fetal brain magnetic resonance imaging (MRI), resolution of the ventriculomegaly by the third trimester, and no abnormalities on newborn cranial ultrasound. One fetus had an isolated increased nuchal translucency measurement of 4.5mm that resolved by the second trimester. The third fetus with an ultrasound abnormality had isolated cleft lip and palate. All fetuses in the amniocentesis group had normal molecular karyotypes (Table 1).

### Amniotic fluid EVs in the second trimester and term differ in size and biogenesis

Western blotting confirmed the presence of EV-enriched protein markers Alix, CD9 and CD63 in the isolated EV samples (Fig 1A). Alix is a protein associated with the EV biogenesis pathway (ESCRT), while CD9 and CD63 are tetraspanins (22, 26). The relative quantities of Alix and CD9 (normalised to the total blot stain in Supplementary Fig 1A) were significantly higher in the term AF-EVs (Fig 1B). Complete blots for Alix, CD63 and CD9 are in supplementary figures 1B and 1C. The enrichment of Alix suggests a greater fraction of endosomal origin EVs (exosomes) in term AF-EVs, compared to that of the second trimester.

**Figure 1.**
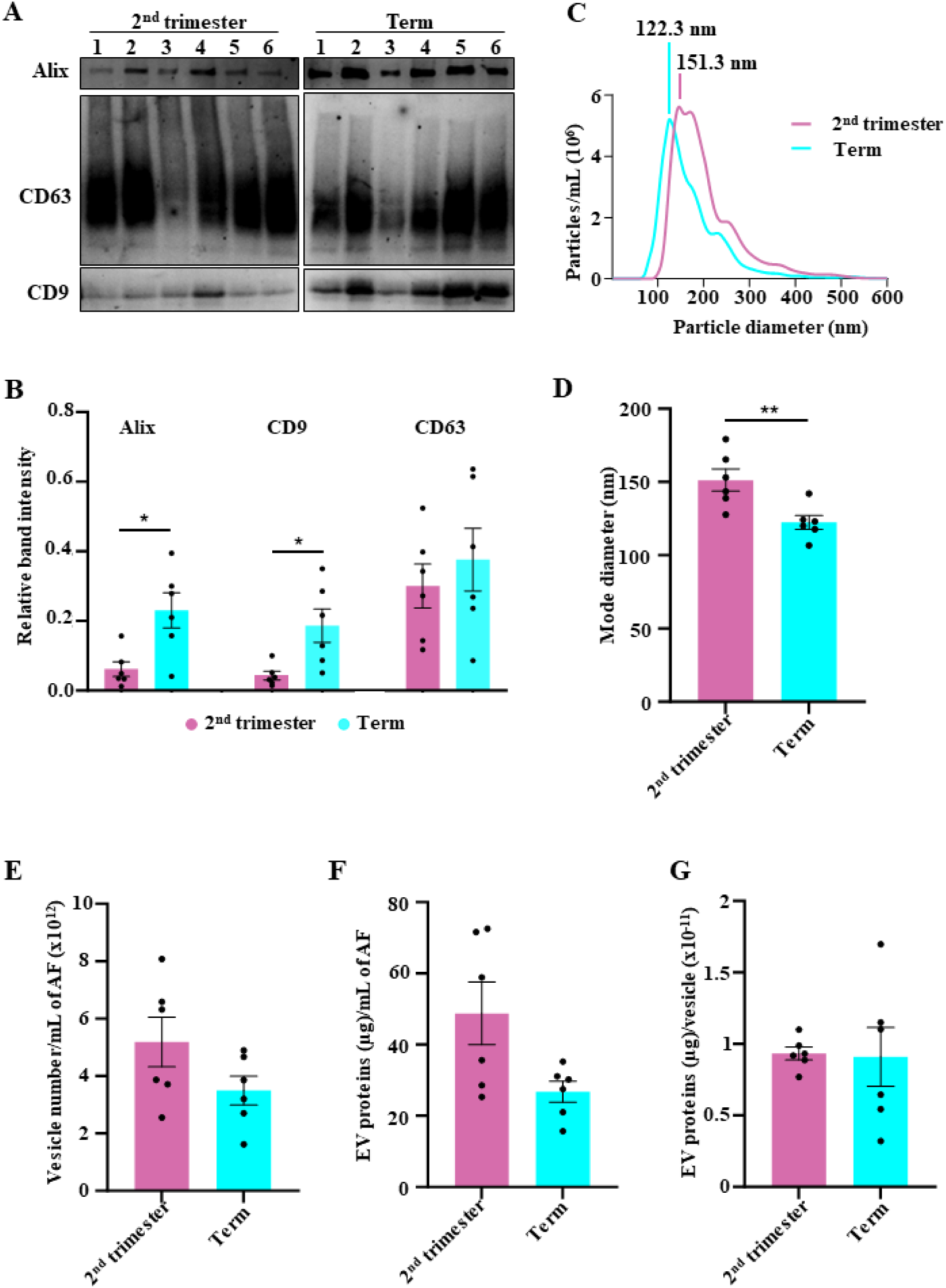
Amniotic fluid EVs at term are smaller in diameter compared to second trimester. (A) AF-EVs were characterised using Western blotting for EV enriched markers Alix, CD9 and CD63. (B) Densitometric analysis of the Western blot confirmed the enrichment of Alix and CD9 in term AF-EVs. (C) Size distribution of AF-EVs was analysed by nanoparticle tracking analysis. The graphs indicate (D) the mode diameter of the AF-EVs, (E) AF-EV concentration, (F) EV protein concentration in AF, and (G) EV protein amount (µg) per vesicle. Statistical analysis used the Students’ T-test with a confidence interval of 95%, in GraphPad Prism 10.1.0 (316). Error bars represent Mean +/- SEM. *<0.05 **<0.01. AF-EVs: Amniotic fluid-extracellular vesicles.

The mode diameter identified by nanoparticle tracking analysis for the second trimester (151.3 nm) was significantly higher compared to that of term AF-EVs (122.3 nm) (Fig 1C and 1D). The EV concentration of AF stayed unchanged between the two gestations (Fig 1E).

Quantification of EV proteins revealed that neither the EV protein concentration in AF (Fig 1F) nor the protein amount (µg) per vesicle (Fig 1G) changed according to gestation.

### Amniotic fluid EVs have unconventional morphologies

We visualised three AF-EV samples from each group, using cryo-electron microscopy and identified different morphologies. Most EVs were simple, with the conventional morphology of single-membrane bound round (orange arrows) or elongated shapes (green arrows) (Fig 2A). EV corona was visible on some vesicles (pink arrows) (Fig 2A). This layer on the outside of the EV membrane represents the collection of proteins, lipids, carbohydrates and nucleic acids a vesicle acquires from its environment (22, 27, 28). While some EVs had multiple membranes (blue arrows), others had multiple compartments (purple arrows) (Fig 2B). We also observed multiple EVs attached with presumed membrane proteins (yellow arrows) (Fig 2C).

**Figure 2.**
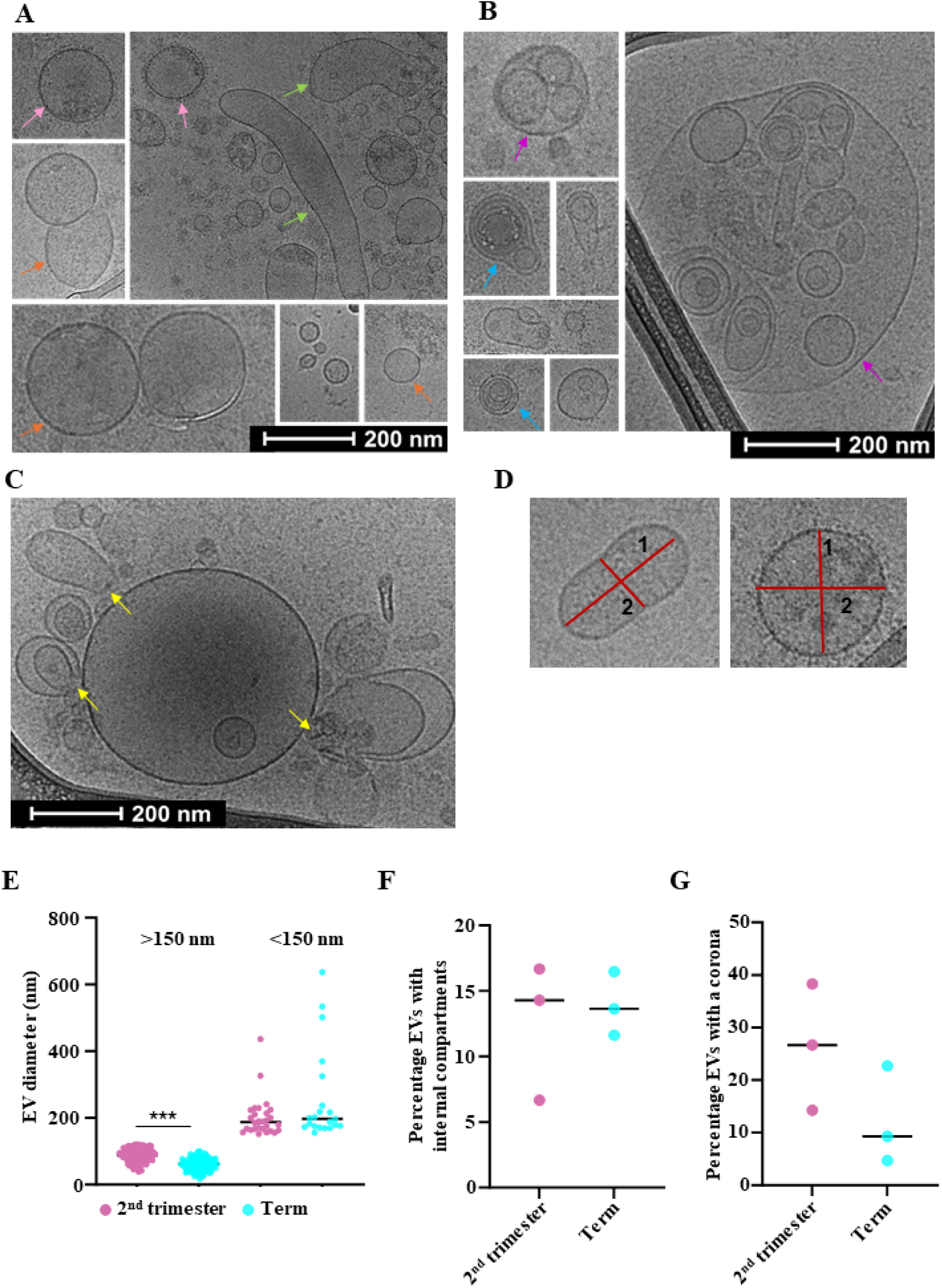
Amniotic fluid EVs have different morphologies. Cryo-electron microscopy identified vesicles with (A) single-membrane bound round (orange arrows) or elongated (green arrows) shape, vesicles with corona (pink arrows), (B) multiple membranes (blue arrows) and/or multiple compartments (purple arrow). (C) Multiple EVs attached together with protein-like structures (yellow arrows) were also visible. (D) EVs were measured in 2 perpendicular axes using ImageJ. Strip charts represent (E) the number of vesicles smaller and larger than 150 nm (each data point denotes a vesicle), (F) the percentage of vesicles with one or more internal compartments and (G) the number of vesicles with a corona, at each gestation. Statistical analysis used the Man-Whitney U test (E) and the Students’ T-test (F, G), with a confidence interval of 95%, in GraphPad Prism 10.1.0 (316). ***<0.001.

To estimate their size, we measured EVs in 2 perpendicular axes (Fig 2D). Since averaging out the two measurements would misinterpret the elongated EV morphology, we used the larger measurement for the analysis. Figure 2E represents the number of EVs for each gestation, separated into two categories based on size, smaller or larger than 150 nm. The significant difference in the number of EVs smaller than 150 nm between gestations complemented the pattern observed in nanoparticle tracking analysis (Fig 1C and 1D).

We manually curated and counted the EVs with different features. Approximately 15% of all EVs had single or multiple compartments internal to the lipid bilayer membrane (Fig 2F). Approximately 25% of 2^nd^ trimester and 10% of term EVs had a corona (Fig 2G).

### Amniotic fluid EV proteome varies in a gestation-dependent manner

We employed label-free proteomics to analyse the EV proteome. Using the DIA type analysis, we identified a total of 4137 proteins (Fig 3A). There were 64 proteins uniquely expressed (present in all 6 replicates) in the second trimester and 13 in the term AF-EVs. Tables 2, 3, and Supplementary Table 1 indicate these proteins and their potential involvement in fetal growth. Some common functions implicated with second-trimester enriched proteins were cell proliferation, differentiation, angiogenesis and neurodevelopment. These proteins were also involved in cell signalling pathways associated with development, such as MAPK, PI3K/Akt, NF-kB and Wnt (Table 2 and Supplementary Table 1).

**Figure 3.**
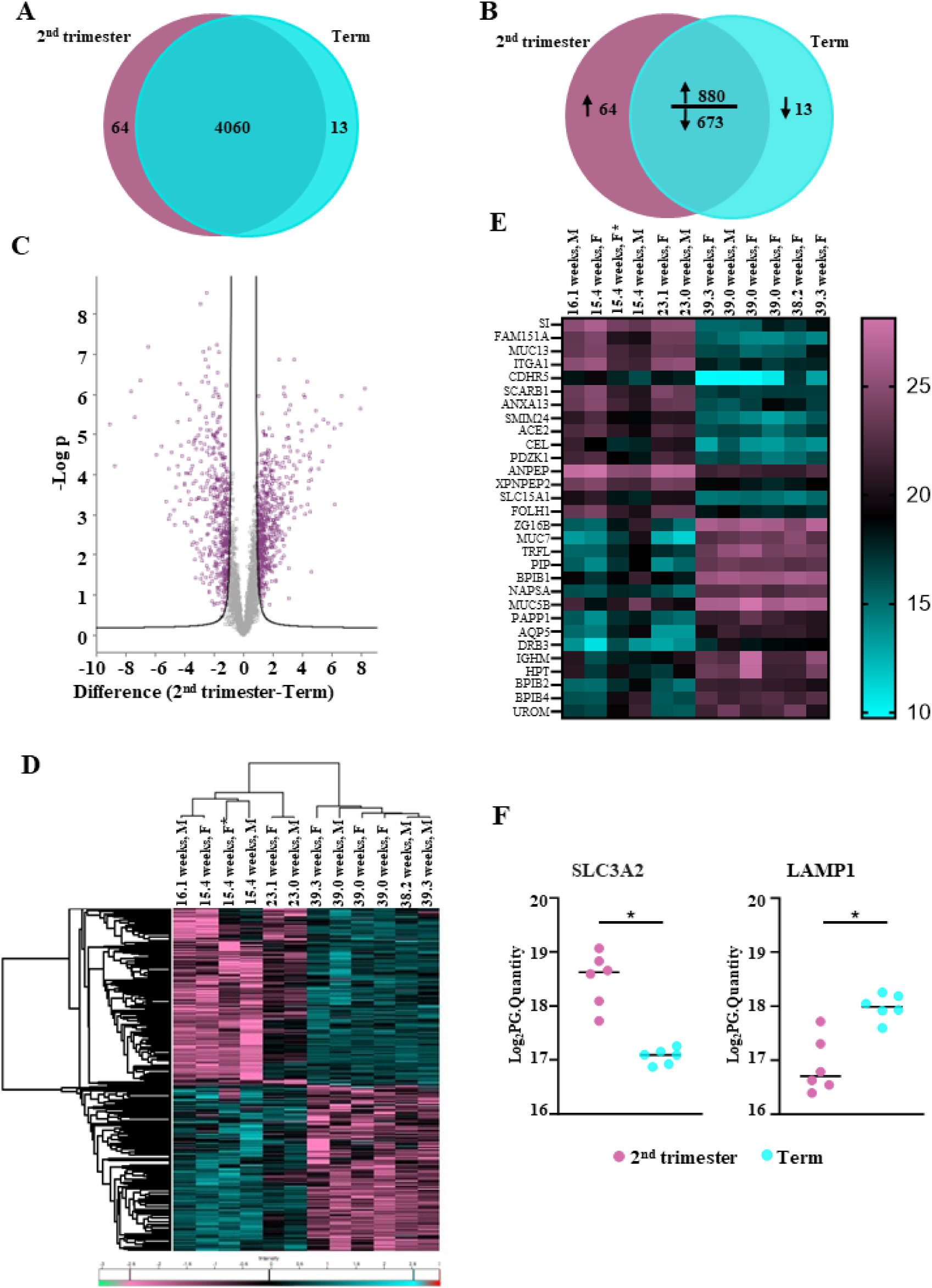
Amniotic fluid EV proteome changes according to gestation. (A) Tandem mass spectrometry identified a total of 4137 proteins, with 64 proteins unique to the second trimester and 13 proteins unique to term AF-EVs. (B) 880 proteins were upregulated, and 673 were downregulated in second-trimester AF-EVs compared to term ones. (C) A total of 1090 proteins were significantly dysregulated between the two gestations (FDR:0.05, S_0_:2). Heatmaps represent (D) the normalised log_2_ transformed values for the significantly dysregulated proteins and (E) the 30 most statistically significant differentially expressed proteins. (F) Ectosome-specific marker SLC3A2 was significantly abundant in second-trimester derived AF-EVs, while exosome-specific marker LAMP1 showed a significant increase in abundance in term-derived AF-EVs. Statistical tests were performed on Perseus using unpaired, 2-tailed Student’s T-test with FDR of 0.05 and 250 randomisations. *<0.05

**Table 2.** The physiological role of the most abundant 15 proteins exclusive to the second-trimester group.

| Gene symbol | Biological Pathways and potential role in human growth and development |
| --- | --- |
| RBP2 | Retinoic acid signalling, Uptake and metabolism of retinoids (29), Potential role in enteric nervous system development (30) |
| TM4SF20 | Cell motility and adhesion (31), Influence neural and craniofacial development (32) |
| ASAH2 | Modulation of PI3K/Akt signalling pathway through ceramide metabolism (33), |
|  | Regulates neuronal differentiation and cell cycle (34) |
| REG4 | Cell proliferation and differentiation (35), Particularly in the gastrointestinal tract (36) |
| ADGRG7 | G protein-coupled receptor activity (37), Potential critical function in angiogenesis, organogenesis, neurodevelopment, and myelination (38) |
| UNC93A | May have a role in sensing nutrient limitation, cellular energy levels and autophagy, Mammalian expression in brain and peripheral tissues (39) |
| SERPINI2 | Involved in insulin sensitivity, fibrinolysis, and inflammatory response (40), Potential role in pathways controlling eye growth and development (41), Serpin-loaded EVs used to promote wound healing in mice (42) |
| SLC17A2 | Transport of organic anions, Involved in vesicular storage of the neurotransmitter and urate metabolism (43) |
| SULT1E1 | Estrogen homeostasis (44), Regulation of hormonal signalling pathways during fetal neurodevelopment (45) |
| IL18R1 | Crucial role in immune system development (46), Angiogenesis (47) |
| NUDT12 | Regulation of nucleotide ratios according to changing conditions (48), Mutations may cause intellectual disability and polyneuropathy (49) |
| GRM2 | Glutamate receptor signalling, Regulates MAPK/ERK1/2 signalling pathway, Mediates neurotransmission and influences the development and integration of neurons (50) |
| GLTPD2 | Involved in protein glycolipid transfer (51), Upregulated during necrotising enterocolitis (52) |
| NLRP6 | Regulates caspase-1, NF-kB and MAPK signalling (53, 54), Inflammasome formation and host defence against microorganisms (55) |
| CELA3B | A serine protease secreted by the pancreas (56) |

*This table summarises the 15 most abundant of the 64 unique proteins in the second trimester and* *their potential role in human physiology, growth, and development.*

**Table 3.** The physiological role of the 13 proteins exclusive to the term gestation group.

| **Gene symbol** | **Biological Pathways and potential role in human growth and development** |
| --- | --- |
| BPIFA2 | Involved in innate immunity, antibacterial defence (57, 58), salivary defence, and surfactant properties crucial for oral health (59) |
| TMEM52B | Regulates cell growth, proliferation and differentiation through EGFR signalling pathway (60), High expression in the kidney (61), Mutations may result in congenital heart defects, orofacial clefts and neural tube defects (60) |
| HLA-DPA1 | Antigen presentation and immune response (62), Potential role in maternal-fetal immune tolerance (63) |
| GKN1 | Exclusively and abundantly made in the stomach. Gastric mucosal defence and homeostasis (64, 65) |
| SLC39A8 | Congenital mutations may result in abnormalities including stunted growth, smaller sized liver, kidney, lung, and cerebrum (66) |
| SUCNR1 | Metabolic sensing, inflammation (1), Haematopoiesis in the bone marrow (67) Influences placental function (68), lipolysis inhibition in adipose tissue (69) |
| MIA3 | Protein trafficking (70), Collagen synthesis (71), Important for extracellular matrix organisation, impacting fetal tissue development |
| PTPN21 | Regulates cell adhesion and migration (72), and signalling pathways crucial for cellular growth and differentiation (73) |
| CAVIN3 | Caveolae formation (74), Regulation of muscle cell function and development, |
|  | potentially impacts cardiovascular development (75) |
| MENT | Chromatin organisation, structure and gene expression regulation (76) |
| HHIP | Hedgehog signalling pathway (77), Essential for tissue patterning and lung development (78) |
| SNAPC4 | Transcription regulation (79) |
| ARHGAP29 | Involved in Rho GTPase signalling (80), Important for cell shape and movement, craniofacial development (81) |

*This table summarises the EV proteins unique to the term gestation (13) and their potential role in* *human physiology, growth and development.*

At term, the unique proteins were primarily responsible for immunity and defence, while some proteins corresponded to the regulation of cell proliferation and tissue patterning, implying controlled growth (Table 3). From commonly expressed proteins, 880 were highly abundant, and 673 were sparsely expressed in the second-trimester AF-EVs compared to term (Fig 3B).

We identified 1099 proteins that were significantly differentially expressed between the two gestations (Fig 3C) (false discovery rate of 0.05 and S0=2). Figure 3D shows the heatmap with unsupervised hierarchical clustering of significantly dysregulated proteins. The proteins are arranged into two distinct main hierarchical clusters in this heatmap, indicating the clear separation of these proteins between the two gestations. The protein signatures of the first four samples corresponding to early second trimester (15-16 weeks) clustered more closely. The clustering gradually changed towards the fifth and sixth samples, representing late second-trimester gestations (23 weeks), before the signature completely changed in the term samples (38-39 weeks).

The group of 30 proteins with the highest fold changes are indicated in Figure 3E. The first 15 proteins were highly abundant in the second trimester derived AF-EVs and were identified with terms such as carbohydrate metabolism (SI) (82), cell adhesion (ITGA1) (83) and CDHR5) (84), lipid transport (SCARB1) (85) and peptide transport (SLC15A1) (86). The following 15 proteins were highly abundant in term-derived AF-EVs. Many of them, such as PIP (87), UMOD (88), HLA-DRB3 (89), BPIFB4 (90) and IGHM (91) were implicated in the immune response. MUC7 is mainly present in saliva and the respiratory tract (92), while AQP2 contributes to saliva secretion (93).

The results of the proteomic analysis showed significant enrichment of the ectosome-exclusive marker SLC3A2 (26) in the second-trimester AF-EVs. LAMP1 (26), a protein marker exclusive to exosomes, was significantly enriched in the term-derived AF-EVs (Fig 3F). These findings complement the relative abundance of Alix in the term AF-EVs observed above, supporting the enrichment of the exosome fraction in the term (Fig 1A and 1B).

### Amniotic fluid EV proteome is a representation of fetal growth and maturation

We subsequently performed an enrichment analysis on the significantly abundant proteins identified in each gestation. Proteins enriched in the second-trimester AF-EVs were matched for expression in organs such as ovary, pancreas, brain, liver and bone marrow (Fig 4A). AF-EV proteome at term demonstrated a significant similarity to other human body fluids such as saliva, tears, cervicovaginal fluid, and gastric juice (Fig 4B). Both gestations indicated a significant enrichment of placental and urine proteins.

**Figure 4.**
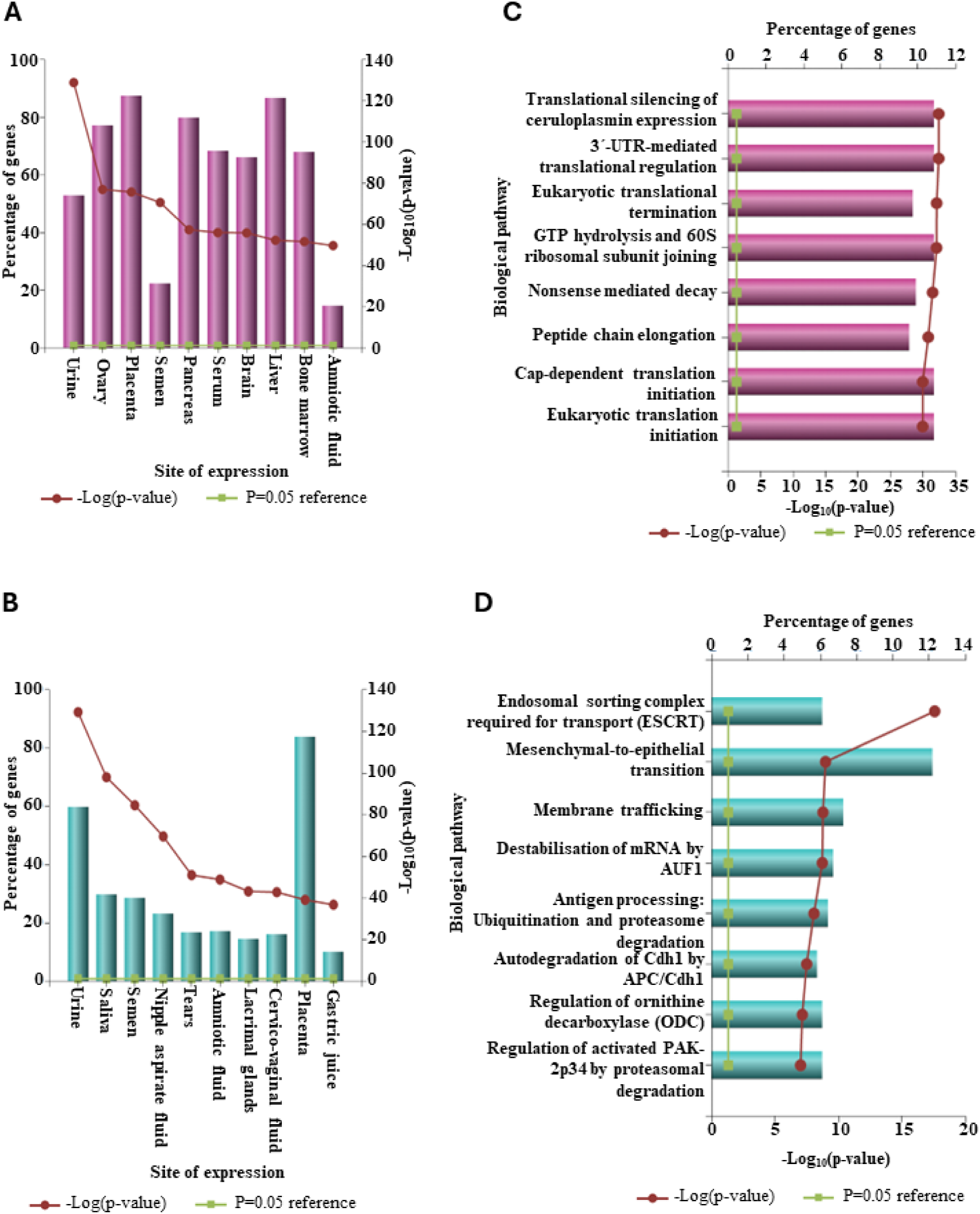
Amniotic fluid EV proteome represents gestational age-dependent fetal growth and development. Enrichment analysis (Funrich) of proteins uniquely expressed and significantly abundant in (A) second trimester and (B) term were matched with distinct site-specific protein signatures using UniProt, HPRD, Human Protein Atlas, Human Proteome Browser, Human Proteome Map, ProteomicsDB and Human Proteinpedia as background data resources (24). A similar analysis was conducted to identify biological pathways enriched in (C) the second trimester and (D) term, using a combination of databases including Reactome, National Cancer Institute Data Catalog, Cell map, HumanCyc, and NCI-Nature Pathway Interaction Database (25).

The enrichment analysis for biological pathways matched second-trimester-enriched AF-EV proteins with translation regulation, ribosome activity, and protein synthesis-related activities (Fig 4C), indicating the development the fetus undergoes during the second trimester.

Proteins enriched in the term AF-EVs showed a significant involvement with biological processes that are important for bringing the fetal development phase to a halt (Fig 4D). Epithelial-to mesenchymal-transition (EMT) is a cellular signalling pathway instrumental in morphogenesis and organogenesis during fetal development (94, 95). The reverse process, Mesenchymal-to-epithelial transition (MET), helps to generate somatic cells (96). We observed the activation of the MET pathway at term gestation. APC/Cdh1 complex, ornithine decarboxylase and active PAK2p34 all play essential roles in mitosis (97), cell growth and differentiation (98) and cell survival (99), respectively. The regulation of these proteins towards term gestation indicates the easing of the fetal development phase. Moreover, adaptive immunity activation towards the end of gestation may be implied in “antigen processing via ubiquitination and proteasomal degradation” as the fetus approaches birth.

We detected a significantly high number of proteins associated with the Endosomal Sorting Complex Required for Transport (ESCRT) machinery in term AF-EVs. This is a signalling pathway and the critical regulator of multivesicular body (MVB) formation and protein sorting into EVs (100, 101). This observation explains the enrichment of exosomes in the term AF-EV pellet compared to that of the second trimester, as observed in Figures 1A, 1B, and 3F.

### Proteomics results after excluding the fetus with a congenital anomaly

Due to the presence of one sample with a congenital anomaly, we repeated the proteomics analysis for five pairs after excluding the fetus with the bilateral cleft/palate and the matched control. We observed a similar overall expression profile, with nine of the ten protein enrichment sites overlapping in the second trimester (Supplementary Fig 2A and Fig 4A) and term (Supplementary Fig 2B and Fig 2B). Similarly, the biological pathways implicated in the second-trimester (Fig 4C and Supplementary Fig 2C) and term (Figure 4D and Supplementary Figure 2D) enriched proteins remained unchanged between original and smaller cohorts. Therefore, we concluded that retaining these samples in our primary analysis was reasonable.

## Discussion

This study presents strong evidence that EV biogenesis and secretion in the fetoplacental unit experience notable changes throughout gestation, reflecting the physiological adaptation to the requirements of the developing fetus. By isolating and characterising the small EVs fraction, comprising both exosomes and small ectosomes, we observed a distinct shift in the predominant EV subtype between the second trimester and term pregnancies. Specifically, second-trimester AF-EVs were enriched in ectosomes, which bud directly from the cell membrane, while term AF-EVs were predominantly exosomes, originating from the endocytic pathway. This shift in EV subtype, correlated with significant differences in vesicle diameter, underscoring the likelihood that these vesicles have unique biological roles at different stages of fetal development. Additionally, the biogenesis and secretion of ectosomes are generally considered less energy-intensive than the formation of exosomes. This distinction arises because ectosome formation bypasses the complex intracellular trafficking and membrane fusion events required for exosome production in the endocytic pathway (102, 103). Similarly, fetal development during the second trimester is characterised by organ development, where the energy consumption is lower compared to the last 8 weeks of pregnancy when the oxygen and nutrient demand increase (104, 105). As such, the fetoplacental unit may prioritise less energy-intensive ectosomes in the second trimester and more energy-demanding exosomes towards term gestation. These findings enhance our understanding of the gestation-dependent characteristics of AF-EVs and open new avenues for exploring their functional implications in fetal physiology.

This study is the first to conduct near-native imaging of AF-EVs with cryo-electron microscopy, revealing their complicated morphologies for the first time. Similar EV phenotypes have been observed in cerebrospinal fluid (106, 107), adipose tissue (108) and neuronal sources (109). Previous research has shown that multiple membranes contribute to the structural integrity of EVs (106); multi-EV clustering may also indicate potential functional microdomains (109). Further research is needed to uncover the biological significance of these different EV phenotypes in fetal development.

We also observed corona on a substantial proportion of AF-EVs. The corona is a recently described phenomenon in EV biology, comprising a complex conglomeration of biomolecules attached to the outer membrane. Coronas form after EV secretion into the extracellular environment and can alter EV functionality and, therefore, contribute to its biological properties (28, 110). Evidence from artificial nanoparticle research demonstrates multiple influences on corona formation, including physical parameters (vesicle diameter, surface curvature and zeta potential) and biochemical parameters (surface proteome) (28, 111, 112). Our results provide the first evidence of this phenomenon in human AF.

Our proteomics analysis revealed the gestation-dependent evolution of the protein cargo in AF-EVs. We observed distinct differences in the protein signatures of second-trimester and term AF-EV samples. There were also detectable differences between the protein cargo of early (15-16 weeks) and late (23 weeks) stages of the second trimester (Fig 3D). Second-trimester AF-EVs showed a high abundance of proteins that drive active cell growth, differentiation, and organ development. In contrast, term AF-EVs contained proteins predominantly responsible for immunity, lung function, and digestion. This shift in protein expression pattern reflects the fetus’s known developmental trajectory across gestation.

In previous studies, our group and others have explored AF cell-free RNA to understand fetal development via a transcriptomic approach (12, 113, 114), revealing similar RNA enrichment patterns to the current study. As EVs are key mediators of extracellular RNA transport (115). This consistency with the AF cell-free RNA literature validates our findings and offers further insights into the biology of AF cell-free nucleic acids.

One sample of the second-trimester group had a congenital anomaly. Given the discovery-driven nature of this study and the similarity of the proteomics findings with and without this sample, we did not exclude it from analyses. However, we acknowledge that some fetuses with and without congenital anomalies may have different AF-EV biology, which provides avenues for further research.

We have shown that AF contains EVs secreted from multiple cell types within the fetoplacental unit. AF and the EVs contained within it circulate throughout the fetal gastrointestinal, respiratory, pulmonary and urinary systems. Additionally, AF is in constant contact with the amniotic membranes and fetal surface of the placenta (11, 116). The presence of EVs in AF and their dynamic biology across gestation suggest important roles in communication between fetal organ systems and between the fetus and the placenta. EVs are capable of docking on distant organs, releasing their biologically active cargo into the target cell’s cytoplasm, thus influencing cellular signalling and metabolic processes of the recipient cell (16, 117). While AF-EVs have been explored as biomarkers for pregnancy complications such as preeclampsia (17), preterm labour (18) and fetal cytomegalovirus infection (118), EV-mediated cellular communication within the fetal compartment remains largely unexplored. Our characterisation of AF-EVs contributes novel insights to this emerging field of research.

Our proteomic analysis uncovered gestation-dependent protein enrichment patterns of AF-EVs, emphasising the importance of considering gestational age when designing studies for biomarker discovery and therapeutic applications. For example, EVs exogenously loaded with Serpin can promote tissue repair in a mouse model of impaired wound healing (42). We showed that second-trimester-derived AF-EVs exclusively express Serpin-2, making this gestation an ideal time for harvesting EVs for this target protein.

Some researchers responded to the restricted availability of amniocentesis samples by harvesting EVs from cultured stem/mesenchymal stromal cells derived from second-trimester AF (119–121)While this is a convenient approach for large-scale applications, it is unknown how the biological properties of EVs of cultured AF cells compare with those derived directly from AF. A comparative analysis of EVs derived from fresh AF versus cultured AF-derived cells may provide further insights into their *in-vivo* biology.

Obtaining a pure population of EVs remains a challenge (122). Each purification consists of EVs originating from more than one subcellular origin, and therefore, our results represent a mixed EV population. Our functional analysis is likely affected by the literature bias inherent in proteomics. Information about fetal development-related molecular mechanisms is sparse (123), and the analysis tools used in this study could not compare the current data set specifically against a fetal proteome data resource. It is also acknowledged that certain pathologies such as cancer, neurodegenerative diseases and cardiovascular diseases are widely researched and thus overrepresented in annotation databases (124, 125). Therefore, in presenting the enrichment analysis, we omitted the results of unapplicable adult pathologies and transformed cell lines, focusing on the findings relevant to human fetal physiology.

### Conclusions

In conclusion, AF-EVs represent an intriguing mechanism for intercellular communication within and potentially outside the fetoplacental unit. This comprehensive investigation demonstrates the diverse morphology and dynamic biology of fetal EV secretion and function. The gestation-dependent EV protein enrichment patterns suggest the purposive nature of EV cargo packaging and the potential of AF-EVs to provide new insights into fetal intercellular communication and physiology. Understanding the biological significance of the different predominant EV subtypes at various gestational stages is a crucial next step in this field.

## Author contributions

L.H. conceptualised the study, acquired funding, supervised the research and provided guidance throughout the project. Contributed to writing, reviewing, and editing and approved the final manuscript. I.A. led the investigation, established protocols, conducted experiments, analysed the data, wrote the original manuscript, reviewed and edited the manuscript. S.B. managed the sample and patient data collection. CS.A. performed the proteomics analysis and contributed to the data interpretation. H.B. optimised cryo-electron microscopy methods. S.R. assisted in conducting experiments and data analysis. B.F. assisted in conducting experiments. N.D.A and N.B. assisted in sample collection. N.H. provided some experimental and software resources. All authors reviewed and provided critical feedback on the manuscript.

## Acknowledgements

We thank the clinical staff in the Department of Perinatal Medicine at the Mercy Hospital for Women, Heidelberg, Victoria, Australia, for their assistance in sample collection. Special thanks to Prof Tu’uhevaha Kaitu’u-Lino and Dr Elif Kadife for providing feedback for this manuscript.

---

**Supplementary figure 1.**
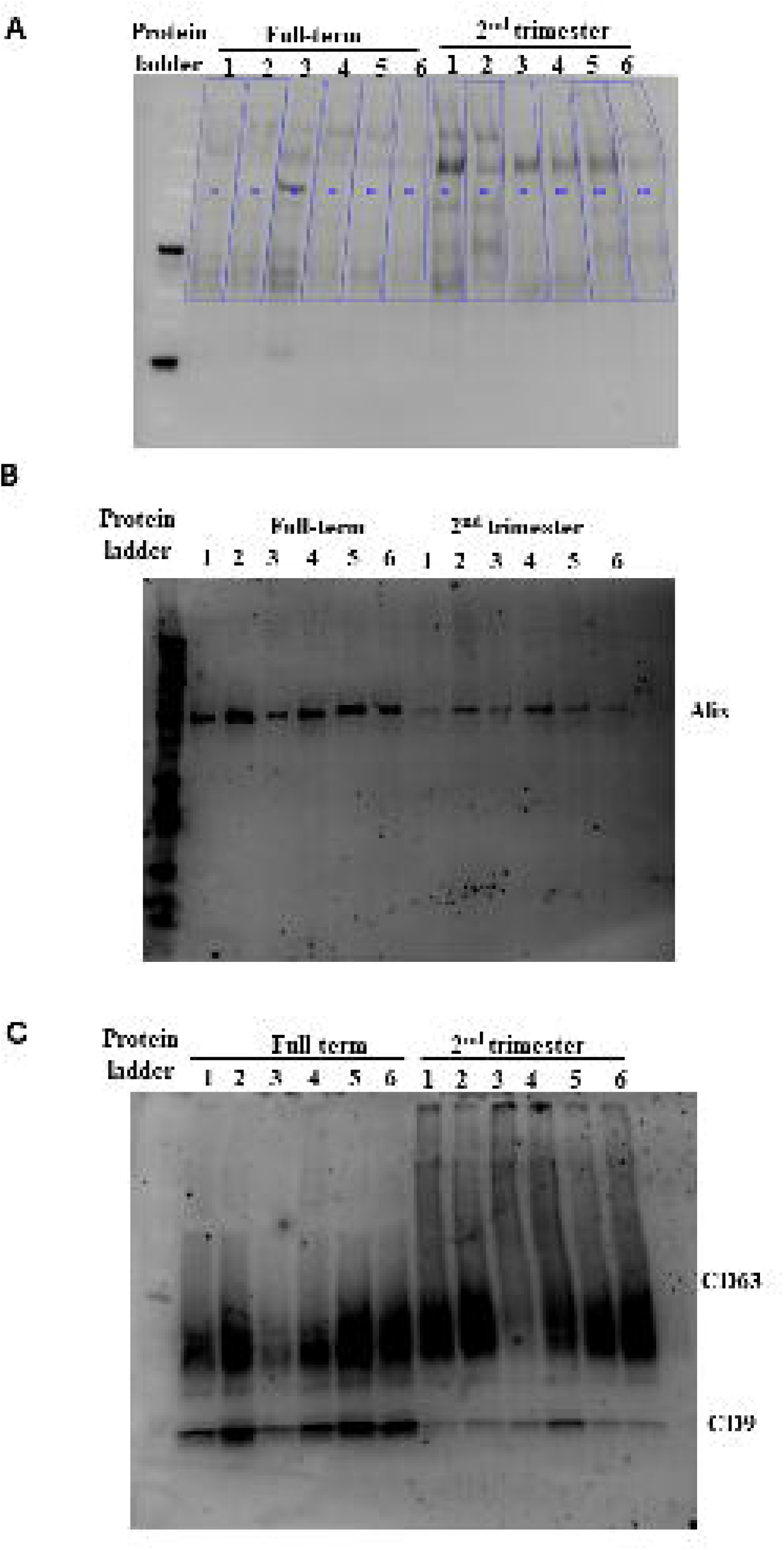

**Supplementary figure 2.**
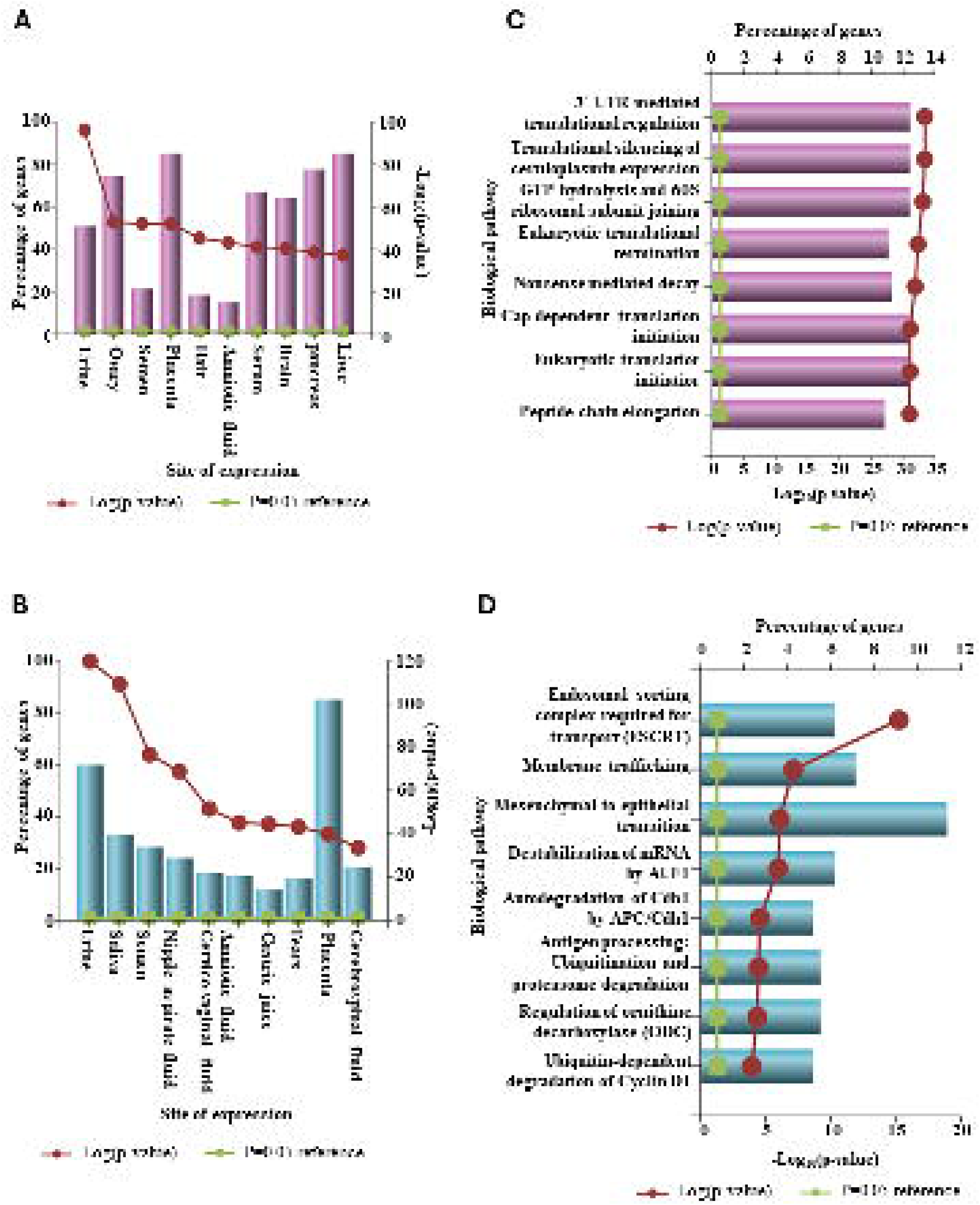

**Supplementary Table 1.**
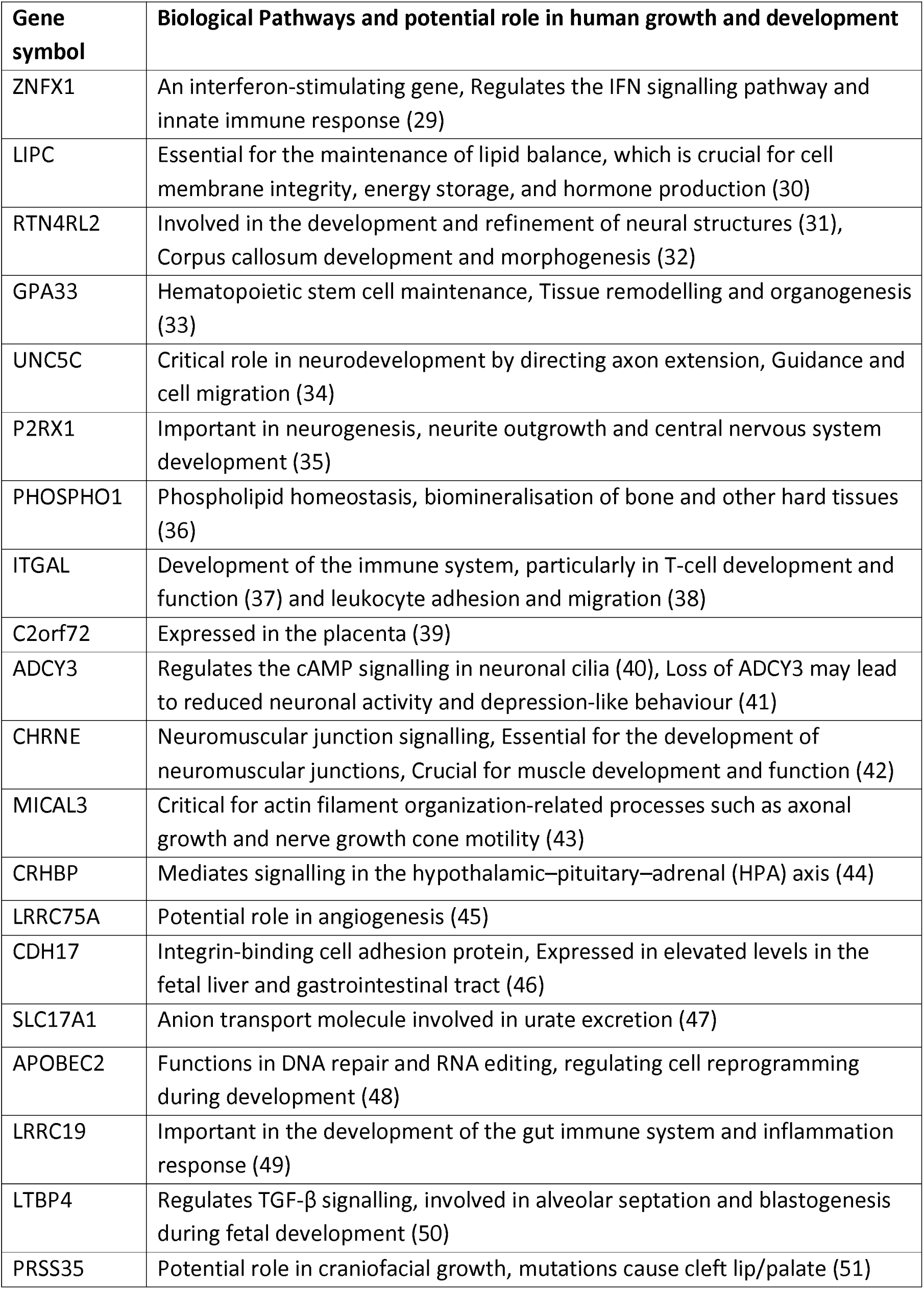
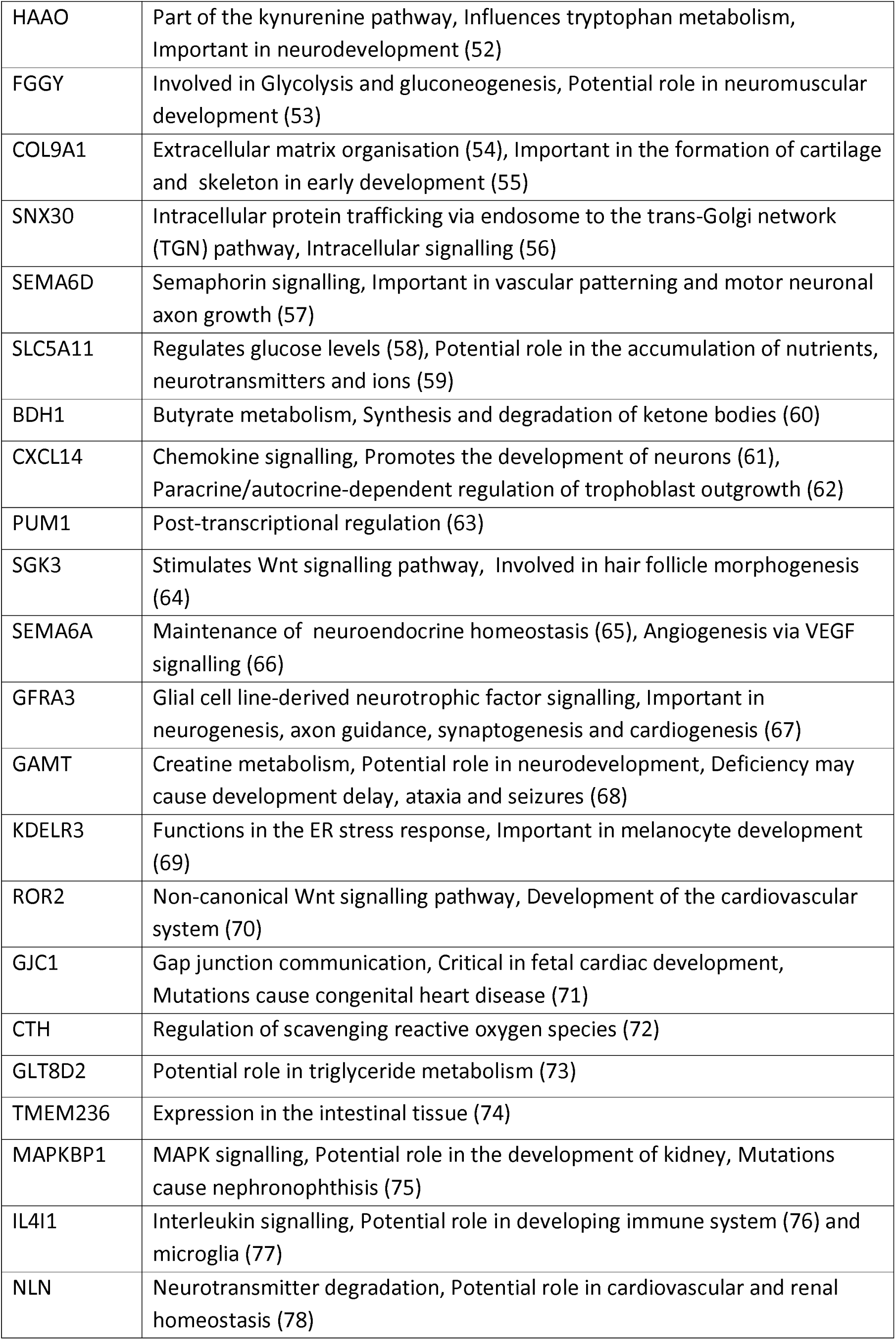
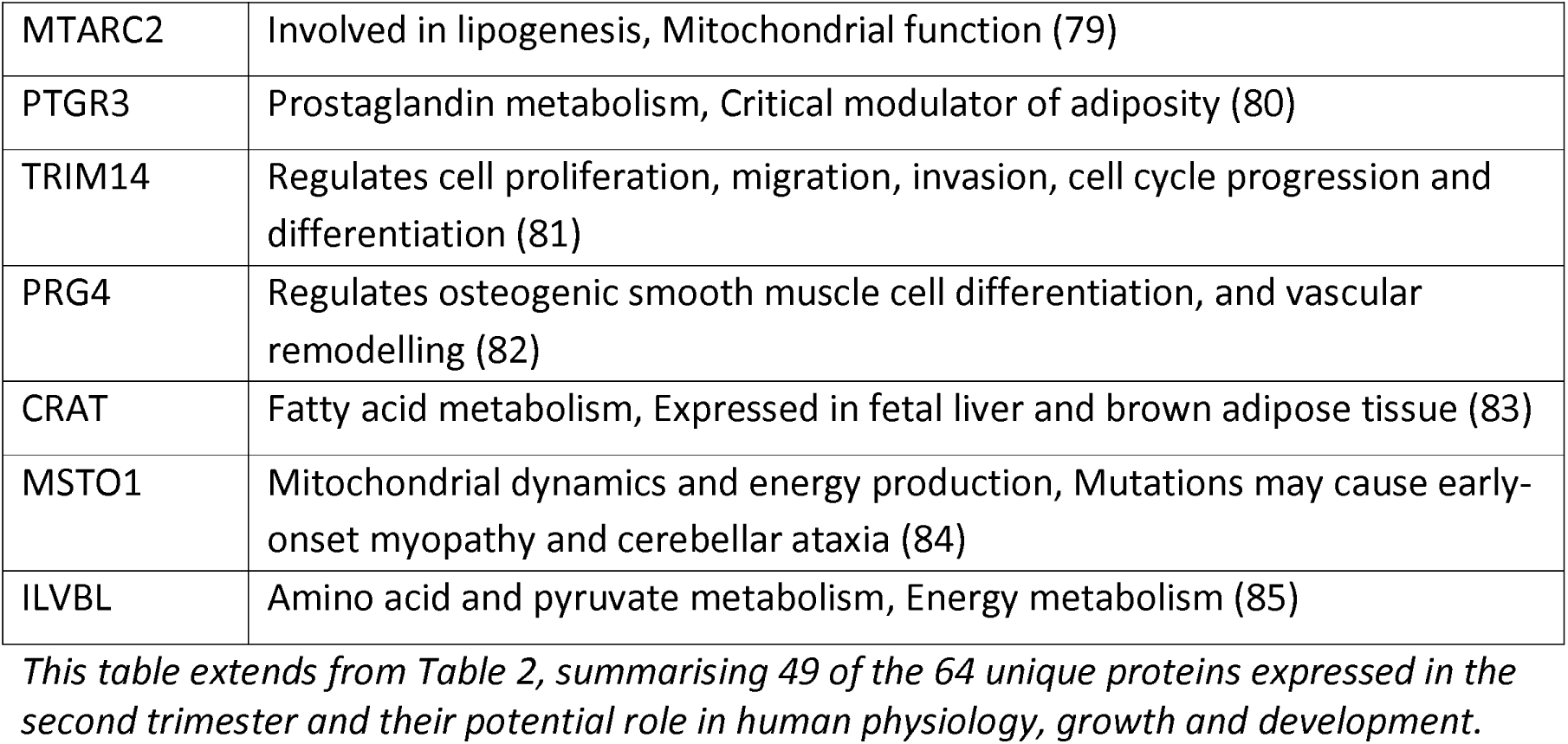
The physiological role of proteins uniquely expressed in the second trimester - an extension to Table 2.

## Notes

http://www.ebi.ac.uk/pride

### Competing Interest Statement

The authors have declared no competing interest.

